# Unconventional ER to Endosomal Trafficking by a Retroviral Protein

**DOI:** 10.1101/2020.07.07.192815

**Authors:** Wendy Kaichun Xu, Yongqiang Gou, Mary M. Lozano, Jaquelin P. Dudley

**Affiliations:** Department of Molecular Biosciences, Center for Infectious Disease, and Institute for Cellular and Molecular Biology, The University of Texas at Austin, Austin, TX

**Keywords:** Rem, mouse mammary tumor virus, endoplasmic reticulum, endosomal trafficking, brefeldin A sensitivity

## Abstract

Mouse mammary tumor virus (MMTV) encodes a Rem precursor protein that specifies both regulatory and accessory functions. Rem is cleaved at the ER membrane into a functional N-terminal signal peptide (SP) and the C-terminus (Rem-CT). Rem-CT lacks a membrane-spanning domain and a known ER retention signal, yet was not detectably secreted into cell supernatants. Inhibition of intracellular trafficking by the drug Brefeldin A (BFA), which interferes with the ER to Golgi secretory pathway, resulted in dramatically reduced intracellular Rem-CT levels. A Rem mutant lacking glycosylation sites was cleaved into SP and Rem-CT, but was insensitive to BFA, suggesting that unglycosylated Rem-CT does not exit the ER or reach a degradative compartment. BFA reduction of Rem-CT levels was not rescued by proteasome or lysosomal inhibitors. Rem-CT has simple glycans, which are necessary for Rem-CT stability and trafficking, but indicate that Rem-CT does not traffic through the Golgi. Analysis of wild-type Rem-CT and its glycosylation mutant by confocal microscopy revealed that both were primarily localized to the ER lumen. A small fraction of wild-type Rem-CT, but not the unglycosylated mutant, were co-localized with Rab5+ endosomes. Expression of a dominant-negative (DN) form of ADP ribosylation factor 1 (Arf1) (T31N) mimicked the effects of BFA by reducing Rem-CT levels, suggesting that Arf1 prevents Rem-CT localization to a degradative compartment. A DN form of the AAA ATPase, p97/VCP, rescued Rem-CT in the presence of BFA or DN Arf1. Thus, Rem-CT uses an unconventional trafficking scheme, perhaps to thwart innate immunity to MMTV infection.

**IMPORTANCE:** Mouse mammary tumor virus is a complex retrovirus that encodes a regulatory/accessory protein, Rem. Rem is a precursor protein that is processed at the endoplasmic reticulum (ER) membrane by signal peptidase. The N-terminal SP eludes ER-associated degradation to traffic to the nucleus and serve a human immunodeficiency virus Rev-like function. In contrast, the function of the C-terminal glycosylated cleavage product (Rem-CT) is unknown. Since localization is critical for protein function, we used multiple methods to localize Rem-CT. Surprisingly, Rem-CT, which lacks a transmembrane domain or an ER retention signal, was detected primarily within the ER and required glycosylation for trafficking to endosomes. Blocking of retrograde trafficking through Arf1 reduced Rem-CT levels, but was not restored by lysosomal or proteasomal inhibitors. The unique trafficking of Rem-CT suggests a novel intracellular trafficking pathway, potentially impacting host anti-viral immunity.

## INTRODUCTION

Mouse mammary tumor virus (MMTV) is the only known murine complex retrovirus with organizational features similar to human pathogenic retroviruses, such as human immunodeficiency virus type 1 (HIV-1) (1-4). Unlike HIV-1, which is difficult to study in its native host, MMTV provides a suitable animal model to elucidate viral interactions within the context of a natural host immune system. Like human complex retroviruses, MMTV also encodes accessory and regulatory genes (4-6). Our studies previously have shown that MMTV encodes both regulatory and accessory proteins from a precursor protein, Rem (2-4). Rem is generated from the doubly spliced MMTV mRNA, which is translated on the endoplasmic reticulum (ER) membrane, and then cleaved by signal peptidase into a 98-amino-acid N-terminal signal peptide (SP) and a 203-amino-acid C-terminal glycosylated protein (Rem-CT). Since Rem is translated in the same reading frame and from the same start codon as the MMTV envelope protein (Env), SP is synthesized from both *rem* and *env* mRNAs, although *env* mRNA is much more abundant (7). After Rem or Env cleavage, MMTV-encoded SP is retrotranslocated from the ER to the cytosol in a p97/VCP ATPase-dependent manner (7, 8). SP then traffics to the nucleus to bind to unspliced MMTV RNA for nuclear export and subsequent steps of virus replication (7-9). Thus, SP has a function similar to human immunodeficiency virus type 1 (HIV-1)-encoded Rev (4). Our recent experiments indicate that Rem and/or Rem-CT act as accessory factors *in vivo* to counteract Apobec-mediated restriction of MMTV replication, particularly to antagonize the mutagenic activity of activation-induced cytidine deaminase (AID) (10). Since protein localization and function are tightly linked, the exact localization and trafficking of Rem-CT within host cells will contribute toward understanding its activity.

Viruses are known to manipulate intracellular membrane compartments to interfere with host antiviral proteins and to promote viral replication (11-21). For example, the mouse cytomegalovirus (MCMV) glycoprotein gp40 arrests the export of MHC I molecules from the ER-Golgi intermediate compartment (ERGIC) to prevent host cell lysis by T cells (11). Poliovirus also blocks trafficking from the ERGIC to the Golgi to enhance its replication (21), whereas vaccinia virus binds host vesicle tethering factors in retrograde transport pathways to assemble new viral particles for replication (20). HIV-encoded Vpu protein prevents the trafficking of the viral receptor (CD4) from the ER to the cell surface, leading to CD4 downregulation, a process that prevents viral superinfection and T-cell activation by antigen-presenting cells (17). Vpu also relocalizes tetherin/BST-2, a protein that interferes with release of many enveloped viruses (12-16, 18, 19, 22-25). The HIV-1 Vpu protein downregulates tetherin from the cell surface, leading to lysosomal degradation or ER-associated degradation (ERAD) and increased viral particle release (12-16, 18, 19). Thus, although retroviruses are known to manipulate the immune function of lymphocytes, MMTV-encoded proteins that subvert cellular trafficking pathways have not been identified.

Proteins synthesized in the ER usually traffic from the ER in COPII-coated vesicles to the ERGIC and then the Golgi network, where they are directed to the cell surface or secreted depending on the presence of transmembrane domains (26, 27). Integral membrane proteins require a sufficiently long transmembrane sequence, which forms an α-helix to span the lipid bilayer (28). ER-synthesized proteins with or without transmembrane domains may be retained within intracellular membrane compartments if they encode retrograde transport sequences, such as a C-terminal KDEL/KKXX for ER retention or, alternatively, an LPYS or KXE/D motif for localization to the Golgi (26, 27). Retroviral Env proteins are synthesized as precursors prior to glycosylation on the surface (SU) portion and folding within the ER. These glycosylated transmembrane proteins must pass a protein quality control system prior to trafficking out of the ER into the Golgi apparatus. The Env precursor is cleaved by the host protease furin within the Golgi into SU and transmembrane (TM) domains that remain covalently linked, and SU glycosylation is further modified (29). Trimers of the SU/TM heterodimers assemble and interact with the Gag precursor for budding (30, 31).

Here we have examined the trafficking of the MMTV Env-related protein, Rem-CT. Our results indicated that Rem-CT is a glycoprotein localized primarily within the ER that lacks a transmembrane domain and a canonical ER retention signal, but is not secreted. In addition, Rem-CT has a unique trafficking scheme from the ER to early endosomes. We observed that Rem-CT intracellular levels were reduced by Brefeldin A (BFA), a fungal metabolite that disrupts protein secretion and collapses the Golgi into the ER (32). Complete rescue of Rem-CT levels after BFA treatment was achieved by expression of a dominant-negative (DN) p97/VCP ATPase or loss of N-linked glycosylation, but not by lysosomal or proteasomal inhibitors. Our data reveal the first virus-encoded protein that employs such a pathway, providing new insights into intracellular protein trafficking and potentially novel antiviral functions.

## RESULTS

### Rem-CT is not a secreted protein

Rem precursor is cleaved by signal peptidase in the ER to generate an N-terminal SP and Rem-CT (Fig. 1A) (7-9). Since Rem-CT is glycosylated and lacks a transmembrane domain or any known canonical KDEL or KKXX sequence at the C-terminus for ER retention (33-35) (Fig. 1A), we predicted that Rem-CT would be secreted into the extracellular medium. To test this idea initially, we transfected 293T cells with a plasmid expressing Rem with an N-terminal GFP tag and a C-terminal T7 tag (GFP-Rem-T7). This approach allowed us to measure SP and Rem-CT levels independently after Rem cleavage by signal peptidase (4, 9). As a positive control, we also transfected cells with an expression plasmid for a secreted form of embryonic alkaline phosphatase, which was tagged at the C-terminus with T7 (SEAP-T7). SEAP is known to be synthesized in the ER and secreted through the classical Golgi pathway (36-38). As expected, both Rem-CT-T7 and SEAP-T7 were detected in cell lysates by Western blotting with T7-specific antibody (Fig. 1B). Rem precursor also was detectable with T7 antibody, and cleaved SP was observed using GFP-specific antibody (Fig. 1B, lanes 1 and 2). To determine if both SEAP and Rem-CT were secreted, supernatants from the transfected cells were collected. Proteins with a mass >15 kDa, which included T7-tagged Rem-CT and SEAP, were concentrated prior to Western blotting using T7-specific antibody. The secretory protein SEAP was easily detectable in cell supernatants (lane 5), whereas Rem-CT was not (lane 7). These results suggested that Rem-CT is not a secreted protein.

**FIG 1.**
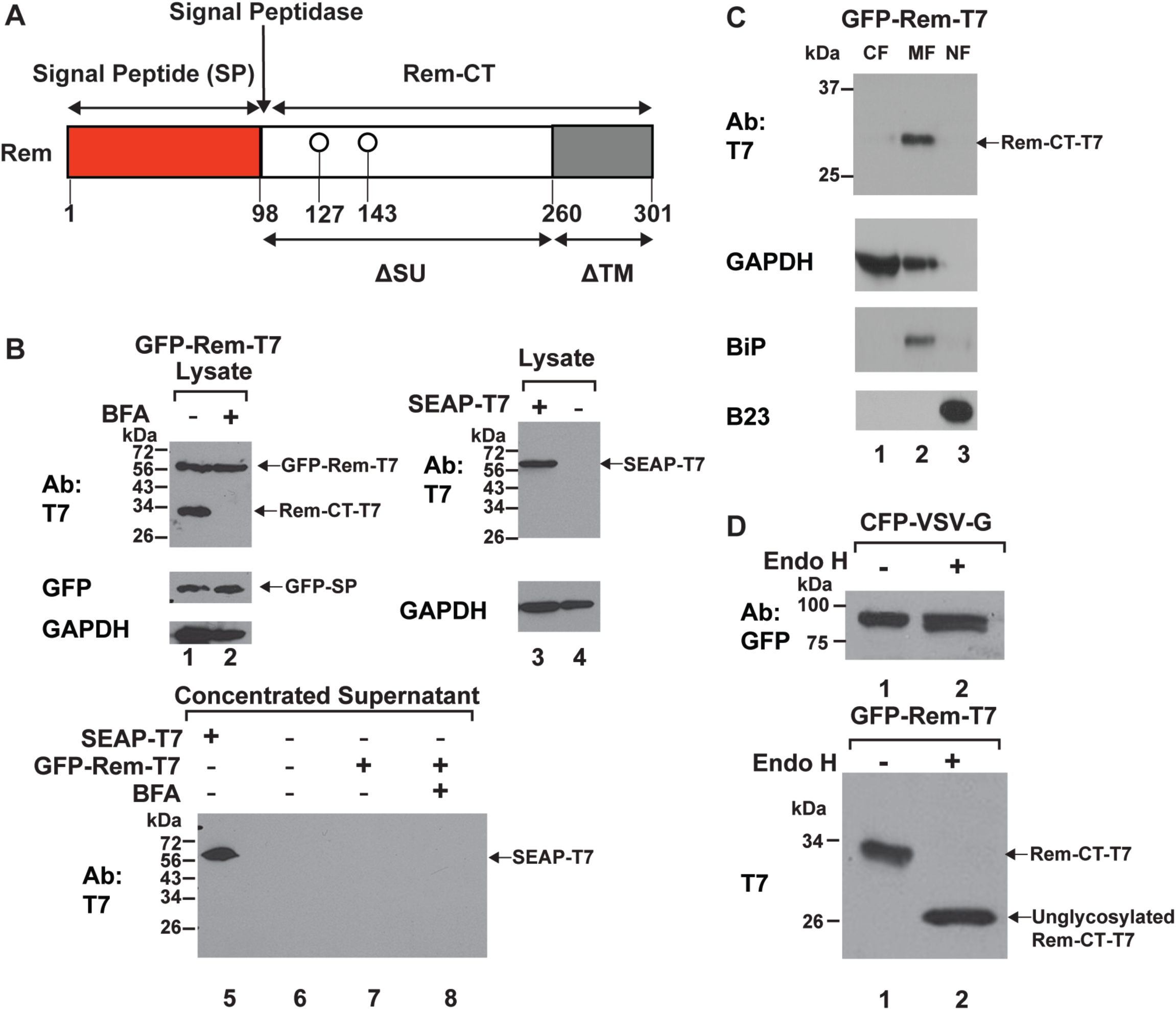
Rem-CT is localized to cytosolic membranes and contains only simple glycans. (A) Diagram of Rem and its cleavage products, SP and Rem-CT. Rem is directed to the ER for translation by the signal peptide sequence. The Rem precursor is then cleaved by signal peptidase at the ER membrane into SP (red box) and Rem-CT (white and grey boxes). The ΔSU and ΔTM designations indicate that Rem represents an in-frame deletion of the Env protein due to a second splicing event within the *env* gene. (B) Rem-CT is not secreted with or without BFA treatment. Cells (293T) were transfected with GFP-Rem-T7 (lanes 1, 2, 7, and 8) or CMV-SEAP-T7 (lanes 3 and 5). In the top panel, cells were treated with or without BFA (lanes 1 and 2). Cells were lysed and subjected to Western blotting with T7, GFP, or GAPDH-specific antibodies. GFP-Rem-T7 is the uncleaved tagged precursor, whereas Rem-CT-T7 is the cleaved C-terminus tagged with T7. The cleaved N-terminus was detected with GFP-specific antibody. Supernatants from transfected cells were collected and concentrated using Amicon® Ultra-15 Centrifugal Filter Units. In the lower panel, concentrated CMV-SEAP-T7 supernatant (50 μg) was loaded as a positive control (lane 5), and concentrated supernatants (500 μg) from cells transfected with the Rem expression plasmid (lanes 7 and 8) were subjected to Western blotting using the T7-specific antibody. Only SEAP-T7 was detected in culture supernatants. (C) Rem-CT localizes to the cytosolic membrane fraction. Cells (293T) were transfected with GFP-Rem-T7 expression plasmid and then lysed and fractionated as described in Materials and Methods. Different fractions were then subjected to Western blotting with T7, BiP, B23, or GAPDH-specific antibodies. (D) Rem-CT glycan is cleavable by Endo H. Cells (293T) were transfected with vectors expressing either CFP-VSV-G (lanes 1 and 2; upper panel) or GFP-Rem-T7 (lanes 1 and 2; lower panel). The indicated cell extracts (20 μg) were exposed to Endo H for 1 h at 37°C prior to Western blotting with GFP- or T7-specific antibodies.

To confirm that Rem-CT is primarily expressed intracellularly, we transfected 293T cells with the GFP-Rem-T7 expression vector and, after 48 h, cells were lysed to obtain cytosolic, membrane, and nuclear fractions (Fig. 1C). As expected, the nuclear fraction was enriched for the nucleolar protein B23 (39, 40), whereas the ER protein BiP (41, 42), was enriched in the membrane fraction. The majority of the enzyme GAPDH was found in the cytosolic fraction, but also was associated with ER membranes as previously described (43-47). Rem-CT showed the same distribution as BiP (lane 2, top panel), whereas we previously have shown that the highest steady state level of SP occurs in nucleoli (4, 7, 9). Therefore, Rem-CT is localized primarily within the intracellular membrane fraction containing the ER protein BiP.

### Brefeldin A reduces intracellular levels of Rem-CT

To confirm the lack of Rem-CT secretion, we used a strategy previously designed to promote the intracellular retention of secreted cytokines (48). This strategy takes advantage of the drug BFA, a fungal metabolite, which disrupts intracellular vesicular trafficking between the ER and Golgi networks (49-54). Specifically, BFA causes dissociation of guanine nucleotide exchange factors (GEFs) specific for ADP-ribosylation factors (Arfs) from the Golgi membrane. Further, BFA inhibits the retrograde transport of proteins from the Golgi to the ER (55-58), and subsequently results in the fusion of ER and Golgi membranes (49, 51, 52). Therefore, BFA inhibits the early steps of protein secretion to intracellularly trap proteins that traffic through the Golgi (32).

To determine the effect of BFA on Rem-CT trafficking, we transfected 293T cells with GFP-Rem-T7. Transfected cells then were treated with 3 µg/ml BFA for 6 h, and lysates from treated and untreated cells were subjected to Western blotting. We anticipated that the presence of BFA would increase the amount of Rem-CT in cell lysates. In contrast, Rem-CT levels were dramatically reduced in lysates from BFA-treated cells, whereas SP and Rem precursor levels were not affected (Fig. 1B, compare lanes 1 and 2). Levels of GAPDH confirmed the integrity of lysates and equal loading of protein in each lane of the gel. In agreement with our failure to detect Rem-CT in concentrated supernatants (Fig. 1B, lanes 7 and 8), results with BFA were consistent with lack of Rem-CT secretion. Together, our data indicate that Rem-CT does not traffic through the classical intracellular pathway involving the Golgi network.

Protein glycosylation is known to be modified after trafficking through the ER and Golgi due to the presence of specific glycosyltransferases (59). Following the addition of simple glycans in the ER, these polysaccharides are modified to a more complex structure by glycosyltransferases within the Golgi (60). Therefore, lysates from 293T cells transfected with GFP-Rem-T7 were treated with Endoglycosidase H (Endo H). Endo H removes simple glycans, which are present on proteins that fail to traffic to the Golgi compartment (61, 62). As a control for proteins that traffic through the Golgi, we also transfected cells with a tagged form of the vesicular stomatitis virus glycoprotein (CFP-VSV-G) (63, 64). As expected, most glycans of the surface-localized VSV-G protein were not cleavable by Endo H (Fig. 1D, compare lanes 1 and 2, upper panel). Surprisingly, the glycans on Rem-CT were cleaved by Endo H (Fig. 1D, compare lanes 1 and 2, lower panel). Since Endo H only cleaves high mannose or hybrid oligosaccharides that have not been processed by Golgi enzymes (61), Rem-CT does not traverse the medial Golgi where the conversion to complex glycans occurs (59, 60).

### Rem-CT susceptibility to BFA is reduced by lack of glycosylation, but not proteasomal or lysosomal inhibition

Our previous results have shown that Rem-CT has two glycosylation sites at positions 127 and 143 relative to the Rem N-terminus (Fig. 1A) (9, 65). Trafficking of viral and host glycoproteins and exit from the ER is known to be affected by glycosylation (59, 66). Therefore, we used expression vectors for GFP-Rem-T7 or its glycosylation-null variant (N127143Q), which contained mutations of both asparagines to glutamines, for transfection of 293T cells. Lysates of transfected cells were analyzed by Western blotting for Rem-CT and SP. As expected, N127143Q migrated faster than wild-type Rem-CT. However, unlike the wild-type protein, the level of the Rem glycosylation mutant did not change in the presence of BFA (Fig. 2A, compare lanes 1 and 2 to lanes 5 and 6). Further, with the same amounts of transfected plasmids, expression of unglycosylated Rem-CT was more stable relative to glycosylated Rem. Incubation of Western blots with GFP-specific antibody indicated that both the glycosylated and unglycosylated Rem proteins were cleaved by signal peptidase. Similar results were observed with HC11 mouse mammary epithelial cells, which are targets of MMTV infection (1) (Fig. S1). Therefore, BFA effects on Rem-CT are not cell type-specific, but glycosylation is required for the drug to decrease Rem-CT levels.

**FIG 2.**
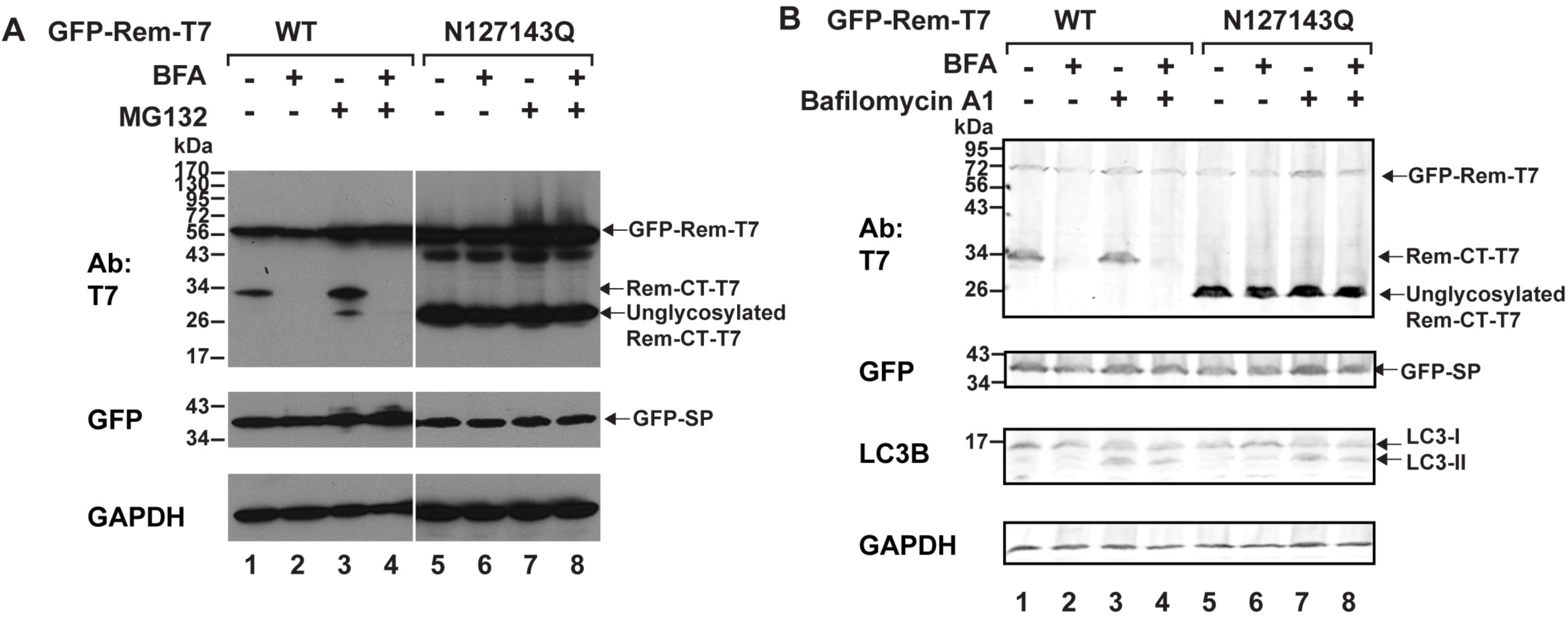
BFA treatment reduces Rem-CT levels and is not rescued by proteasomal or lysosomal inhibitors. (A) Proteasomal inhibitor MG132 did not reverse BFA-mediated reduction of Rem-CT levels. Cells (293) were transfected with equal amounts of expression vectors for GFP-Rem-T7 wild-type (WT) (lanes 1-4) or the N127143Q mutant (lanes 5-8) and treated with BFA (3 μg/ml) and/or MG132 (10 μM). Cell lysates were subjected to Western blotting with T7, GFP or GAPDH-specific antibodies. (B) BFA-induced reduction of Rem-CT levels was not rescued by a lysosomal inhibitor. Transfections were performed as in panel B, and cells were treated with BFA (3 μg/ml) and/or Bafilomycin A1 (1 μM). Cell (293T) lysates were subjected to Western blotting with T7, GFP, LC3B, or GAPDH-specific antibodies. LI-COR imaging was performed after appropriate secondary antibodies were used. The appearance of LC3-II served as a positive control for Bafilomycin A1 activity.

Since the Rem precursor is subject to ERAD, but SP is not, we also tested whether the diminished levels of Rem-CT observed in the presence of BFA were rescued in the presence of the proteasomal inhibitor MG132 (8, 9). As expected, tagged Rem precursor (GFP-Rem-T7) levels increased regardless of the glycosylation status (Fig. 2A, compare lanes 3 and 4 to lanes 1 and 2). SP levels produced by cleavage of the glycosylated Rem protein also slightly increased in the presence of MG132, but SP levels generated from the unglycosylated Rem precursor did not change after proteasomal inhibition. Further, Rem-CT levels generated from the glycosylated Rem protein were not rescued in the presence of BFA by proteasomal inhibition. These results suggest that decreased Rem-CT levels resulting from BFA reorganization of the secretory pathway are not due to ERAD.

Another possibility is that Rem-CT leaves the ER for a secretory compartment and that BFA treatment leads to fusion to lysosomes, which contain large amounts of degradative enzymes (67, 68). Therefore, we transfected 293T cells with expression vectors for either wild-type Rem or the glycosylation mutant N127143Q. A portion of the transfected cells were treated with BFA or BFA plus the inhibitor Bafilomycin A1. This inhibitor blocks the activity of the vacuolar H+ ATPase and prevents the low pH needed for proteolytic enzyme activation within endosomes and lysosomes (69, 70). As anticipated, BFA reduced the intracellular levels of Rem-CT, but not the unglycosylated protein, as detected by Western blots of transfected cell lysates (Fig. 2B, compare lanes 1 and 2 with lanes 5 and 6). Bafilomycin A1 did not rescue wild-type Rem-CT levels observed in the presence of BFA (Fig. 2B, compare lanes 3 and 4) or change levels of N127143Q (lanes 7 and 8). As a control for the activity of Bafilomycin A1, we confirmed the conversion of LC3I to the lipidated form LC3II (71-74). We also isolated exosomes after BFA treatment, but Rem-CT remained undetectable whereas GAPDH can be detected in exosomes (75) as a positive control (Fig. S2). Together, our data suggest that BFA neither induces proteasomal or lysosomal degradation of Rem-CT nor promotes extracellular production of Rem-CT-containing exosomes.

### Rem-CT primarily is localized within the ER

Although our data indicated that Rem-CT is located within intracellular membranes, its exact location has not been determined. Our previous studies using fluorescent microscopy indicated that Rem with a C-terminal GFP tag was not cleaved or folded properly to allow a detectable signal (7) and, therefore, live-cell imaging has not been possible by this method. However, Rem expressing a C-terminal T7 tag (Rem-T7) is cleaved to give functional SP in a dose-dependent manner (Fig. S3). Therefore, we used indirect fluorescence to track Rem-CT within transfected 293 cells. Fluorescence was observed in a mottled pattern within the cytoplasm of transfected cells incubated with primary T7-specific antibody and a fluorescein isothiocyanate (FITC)-tagged secondary antibody (Fig. 3A). This pattern was similar to that obtained after transfection of a plasmid expressing the fluorescent protein mCherry with an N-terminal signal peptide and a C-terminal KDEL to promote ER entry and retention (ER-mCherry) (33, 35, 76). Both Rem-T7 and ER-mCherry were excluded from the nucleus, whereas Rem-CT was highly co-localized with the ER-mCherry (see merged image). As expected, Rem-CT fluorescence was not observed when transfected cells were incubated with BFA (data not shown).

**FIG 3.**
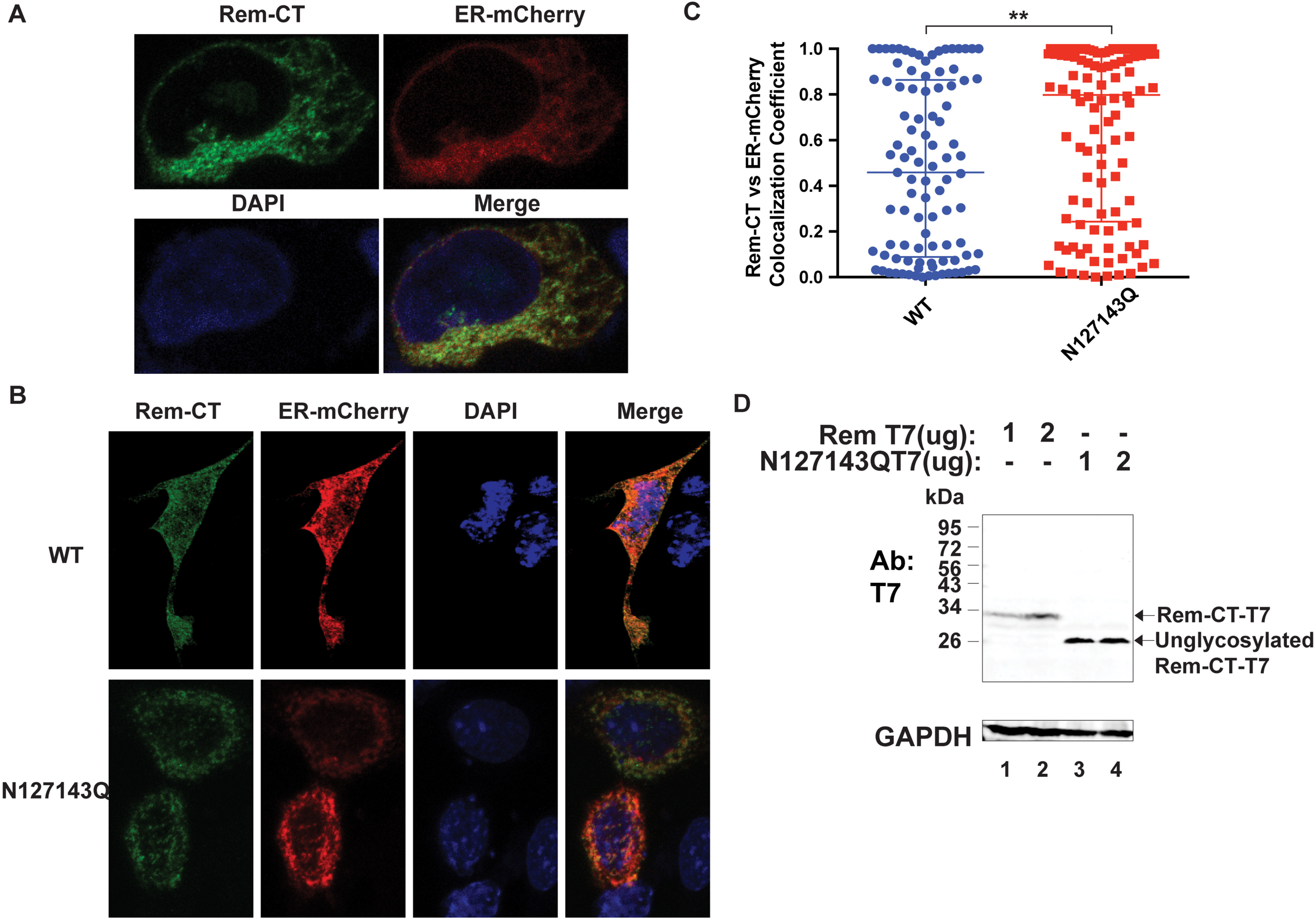
Both wild-type and unglycosylated Rem-CT localize to the ER. (A) Rem-CT colocalized with an ER marker. Cells (293) were co-transfected with expression plasmids for Rem-T7 and ER-mCherry. Rem-CT was detected by a T7-specific antibody followed by Alex Fluor 488-tagged secondary antibody. Images were obtained by confocal microscopy (see Materials and Methods). The green color shows the localization of Rem-CT (upper left) as detected by a T7-specific antibody and an Alex Fluor 488 secondary antibody. The red color detects ER-mCherry (upper right). DAPI stains the nucleus blue (lower left). The lower right panel shows the merged image. (B) Representative confocal microscopy images of wild-type and mutant N127143Q Rem-CT transfected cells. Equal amounts of wild-type or mutant expression constructs were co-transfected into mouse mammary HC11 cells and detected as described in panel A. (C) Scatter plot of Rem-CT wild-type or glycosylation mutant versus ER-mCherry colocalization coefficients. Dots represent colocalization coefficients in individual cells. The middle line in each group represents the median, whereas the highest and lowest lines represent the interquartile range. Either 97 or 108 cells were analyzed for WT and N127143Q, respectively. Statistical analysis was performed using nonparametric Mann-Whitney tests. **p<0.01. (D) Rem precursor is not detectable in mouse mammary cells at the DNA levels used for confocal microscopy. HC11 cells were transfected with the indicated amounts of expression constructs for Rem-T7 WT or N127143Q. Cell lysates were subjected to Western blotting with T7 or GAPDH-specific antibodies. LI-COR imaging was performed after appropriate secondary antibodies were used. Note that the unglycosylated mutant N127143Q shows a faster migration than WT Rem-CT (compare lanes 1 and 2 with 3 and 4).

To further confirm Rem-CT localization, we co-transfected plasmids expressing either Rem-T7 or the N127143Q mutant together with the ER-mCherry expression vector into HC11 mouse mammary epithelial cells (Fig. 3B). Similar to results from 293 cells, we observed Rem-CT colocalization with the ER marker in HC11 cells expressing either the wild-type or the glycosylation mutant (N127143Q). To determine intracellular location more quantitatively, we determined colocalization coefficients, which quantified the ER-colocalized fraction of Rem-CT pixels in multiple individual cells (Fig. 3C). The median value of the colocalization coefficient for wild-type Rem-CT was ∼0.5, whereas the mutant coefficient was significantly higher (∼0.8). As previously reported, higher levels of transfected Rem-T7 expression plasmids led to abundant detection of both uncleaved Rem precursor as well as cleaved Rem-CT (10). Therefore, we performed Western blots of cell lysates from these transfections using T7-specific antibody. The results confirmed that under these conditions only the cleaved Rem-CT was detectable (Fig. 3D). Moreover, similar to results in 293T cells, N127143Q accumulated to higher levels than the wild-type protein. Nevertheless, the distribution of the signals suggests that neither the wild-type nor the glycosylation mutant traffics exactly with the ER-mCherry marker, with the potential to localize elsewhere. These results indicated that the majority of both glycosylated and unglycosylated Rem-T7 remains associated with the ER.

Since Rem-CT lacks a transmembrane domain and a C-terminal KDEL/KKXX retrograde transport signal for ER retention (35, 77, 78), we attempted to map Rem-CT sequences required for its primary localization within the ER. Four deletion mutants were constructed that lacked sequential 50-amino-acid regions starting from residue 103, but C-terminal to the signal peptidase cleavage site (4, 7-10). All mutants retained the C-terminal T7 tag. Expression plasmids for wild-type, deletion mutants, and the N127143Q glycosylation mutant were transfected independently into 293T cells and, after 48 h, cells were lysed for Western blotting with different antibodies. To ensure that the deletions did not affect Rem cleavage in the ER, we examined SP expression with GFP-specific antibody (Fig. 4A and 4B). Compared to wild-type Rem, similar amounts of SP were produced by all mutants with the exception of Δ103-155. Equal loading of protein extracts was confirmed by levels of GAPDH. Although Δ103-155 does not remove the consensus signal peptidase cleavage site, these results suggest that sequences C-terminal to the consensus site affect cleavage efficiency since the level of uncleaved Rem is similar to that observed with the wild-type expression plasmid. To further confirm the reduced generation of SP from the Δ103-155 mutant, we used our quantitative SP reporter assay (4, 7-9). As expected, the wild-type Rem expression plasmid gave ∼12-fold induction of the *Renilla* luciferase reporter at the DNA level transfected relative to the reporter vector alone (Fig. 4C). The Δ103-155 plasmid lacking sequences just downstream of the signal peptidase cleavage site showed a significantly lower SP activity compared to wild-type Rem. Interestingly, our previous results showed that substitution of single glycosylation sites preserved wild-type SP activity, whereas mutation of both glycosylation sites (N127143Q) reduced SP function (9). As expected, the N12713Q mutant also showed a significantly lower SP activity compared to wild-type Rem (Fig. 4C). These results indicated that the majority of the deletion mutants were sufficiently stable to assess Rem-CT levels in the presence and absence of BFA.

**FIG 4.**
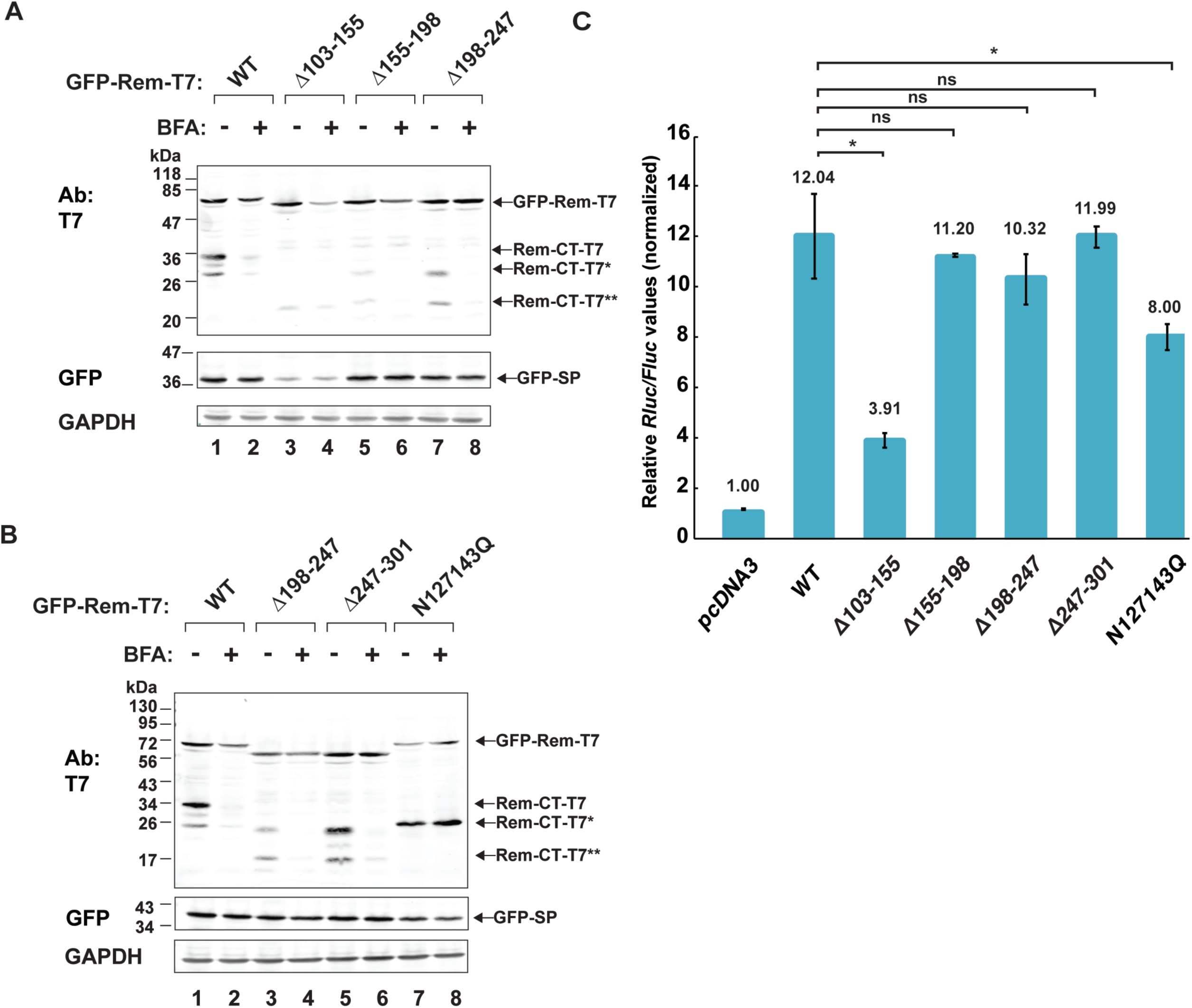
Loss of glycosylation sites rescues Rem-CT levels in the presence of BFA. (A) and (B) BFA sensitivity of Rem-CT deletion and substitution mutants. Expression plasmids for GFP-Rem-T7 wild-type, deletion mutants (GFP-RemΔ103-155-T7, GFP-RemΔ155-198-T7, GFP-RemΔ198-247-T7, and GFP-RemΔ247-301-T7), or the glycosylation site substitution mutant GFP-N127143Q-T7 were transfected into 293T cells and treated with 3 μg/ml BFA as indicated. Whole cell lysates were subjected to Western blotting with T7, GFP, or GAPDH-specific antibodies. LI-COR imaging was performed after incubation with tagged secondary antibodies. The arrows represent different forms of Rem detected by the T7 or GFP-antibodies. GFP-Rem-T7 is the uncleaved precursor, whereas Rem-CT-T7 is the wild-type C-terminal cleavage product. Arrows also point to Rem-CT-T7*, the unglycosylated form of Rem-CT, as well as Rem-CT-T7**, which has a deletion that spans the two glycosylation sites. (C) SP activity of Rem-CT deletion or substitution mutants. Cells (293T) were transfected with 2.5ng of different GFP-Rem-T7 constructs together with Rem-responsive and Rem non-responsive luciferase reporter constructs. Luciferase activities normalized for transfection efficiency were reported as the means and standard deviations of triplicate transfections. The pcDNA3 control sample lacked any Rem expression vector and was set to a relative value of 1.

Half of the transfected cells from the same experiment were treated with BFA prior to analysis using Western blotting with T7-specific antibodies. As anticipated, wild-type Rem-CT levels were greatly reduced in the presence of BFA, and some unglycosylated protein was observed, which co-migrated with the protein produced from the glycosylation mutant N127143Q (Fig. 4B, compare lanes 1 and 7). Also as anticipated, levels of the glycosylation mutant were unchanged in the presence of BFA (Fig. 4B, lanes 7 and 8). The Δ103-155 mutant had low levels of Rem-CT in the presence and absence of BFA, and this protein had a mobility consistent with the deletion and loss of both glycosylation sites. Uncleaved Δ103-155 mutant Rem levels were affected by BFA more than wild-type Rem. Nevertheless, these results were consistent with those from N127143Q, indicating that loss of glycosylation leads to resistance of Rem-CT to BFA. The other mutants Δ155-198, Δ198-247, and Δ247-301 had reduced levels of Rem-CT in the presence of BFA (Figs. 4A and 4B), although only the Δ247-301 mutant had wild-type levels of Rem-CT. Our previous results with N-terminally GFP-tagged Rem mutants containing sequential deletions from the C-terminus also showed that removal of more than 50 amino acids from the Rem C-terminus led to destabilization of SP and uncleaved Rem (7).

Published data indicate that failure to exhibit simple glycans and the correct conformation leads to retention of proteins within the ER, but may also result in ERAD (79, 80). To identify alternative sequences that provide BFA resistance, we also performed a motif search within Rem-CT. These motifs were modified by alanine substitutions, and the subsequent mutants were transfected into 293T cells. Half of the cells were treated with BFA as described for the internal deletion mutants. None of these mutants showed resistance to the drug (Fig. S4). Together, these results suggest that only the loss of glycosylation sites rescues Rem-CT levels in the presence of BFA by promoting retention within the ER.

### Rem-CT traffics to endosomes

Our previous data indicated that Rem-CT is primarily found in the ER and does not traffic through the Golgi typical of other viral glycoproteins. To test whether Rem-CT traffics out of the ER to other cellular compartments, we used confocal microscopy to examine HC11 mammary epithelial cells transfected with plasmids expressing C-terminally T7-tagged wild-type Rem-CT or the glycosylation mutant N127143Q. Transfected cells were then fixed and incubated with T7-specific and Rab5-specific primary antibodies prior to detection with GFP and RFP-tagged secondary antibodies. Rab5 is a regulatory guanosine triphosphatase (GTPase) that serves as a marker for early endosomes (81, 82). We observed co-localization of Rem-CT with Rab5 in a small percentage of transfected cells (Fig. 5A). To quantitate these results, we determined the Rem-CT and Rab5 colocalization coefficient after observation of multiple cells (Fig. 5B). The fraction of wild-type Rem-CT colocalized with Rab5 was very low, indicating that only a small fraction of Rem-CT trafficked to the early endosomes. Alternatively, trafficking of Rem-CT out of the ER to early endosomes may occur very quickly since our results measured only a single fixed time. We did observe a significantly higher colocalization coefficient of Rem-CT with Rab5 compared with that for the glycosylation mutant N127143Q. These data suggest that Rem-CT traffics to the early endosomes, but trafficking is impaired in the absence of glycosylation.

**FIG 5.**
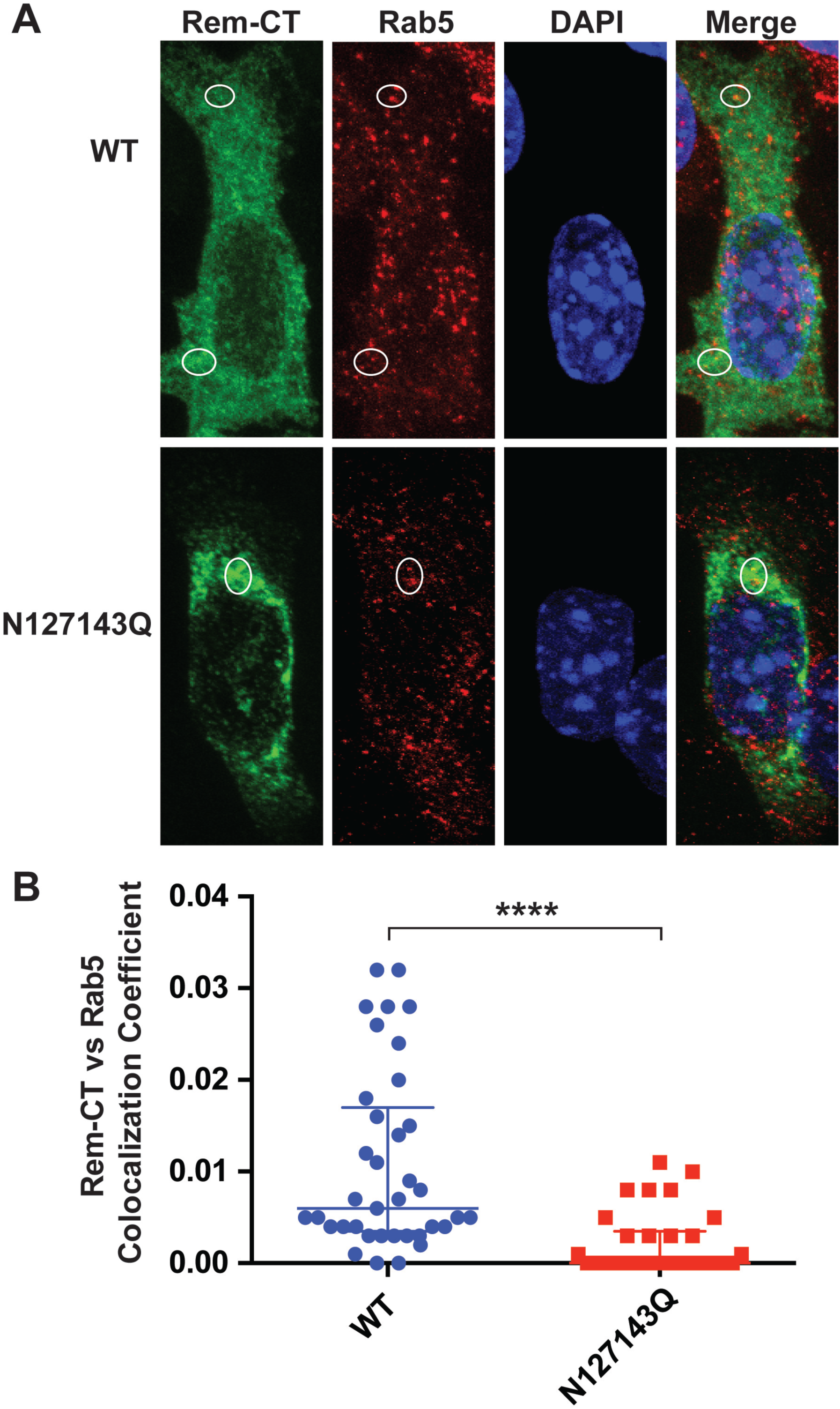
Rem-CT traffics to early endosomes. (A) Representative images of WT and N127143Q Rem-CT stained with the early endosome marker Rab5. Equal amounts of WT and N127143Q Rem-T7 expression constructs were transfected into mouse mammary HC11 cells. Rem-CT was detected using a T7-specific antibody. Appropriate Alex Fluor 488 (green color) and Alex Fluor 594 (red color) secondary antibodies were used for T7 and Rab5, respectively. DAPI (blue) stained the nucleus. Circles mark the colocalization of Rem-CT with Rab5. (B) Quantitation of Rem-CT colocalization with Rab5. Dots represent the colocalization coefficient in individual cells. The middle line in each group shows the median value of the group, whereas the highest and lowest lines (lowest lines are on the x axis) represent the interquartile range. The median values for wild-type Rem-CT and N127143Q were 0.01 and 0.00, respectively. The numbers of cells analyzed were 37 for the wild-type, and 30 for the N127143Q mutant. Statistical analysis performed using nonparametric Mann-Whitney tests indicated a highly significant difference between WT and mutant (****p<0.0001).

Proteins found within early endosomes often traffic to late endosomes prior to lysosomal fusion and degradation (83, 84). Therefore, we also incubated Rem-CT-T7-expressing cells with antibodies specific for the late endosomal marker, Rab7, to determine if Rem-CT is also located in late endosomes. We again observed low levels of colocalization between Rem-CT and Rab7 (Fig. 6A). Quantitation of Rem-CT colocalization with Rab7 in multiple transfected cells was performed to determine the colocalization coefficient (Fig. 6B). Unlike results with Rab5-specific antibodies, these experiments indicated similar low levels of colocalization of wild-type Rem-CT and the mutant N127143Q with Rab7. These data suggest that a portion of both wild-type and glycosylation mutant Rem-CT reach the late endosomes.

**FIG 6.**
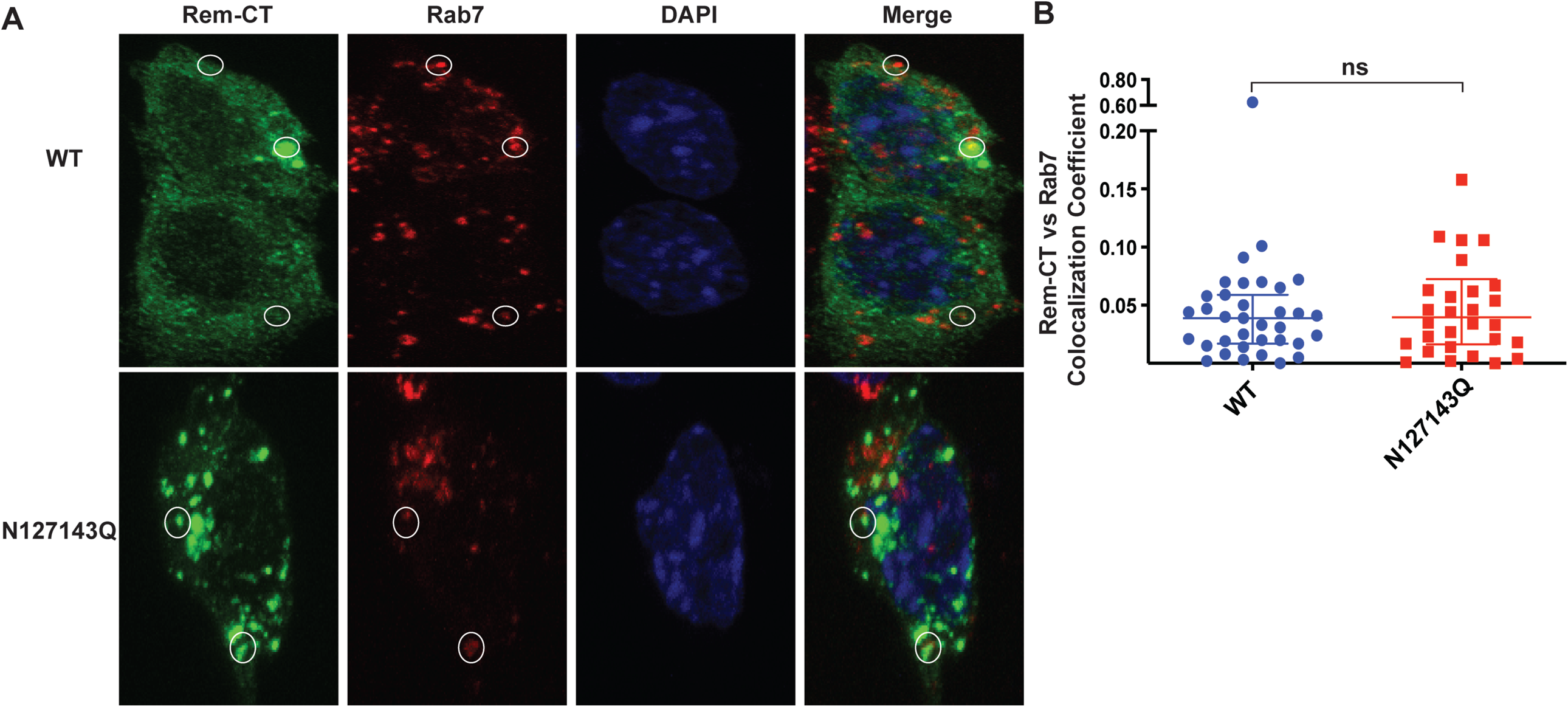
Rem-CT traffics to the late endosomes. (A) Representative images of WT and N127143Q Rem-CT stained with the late endosome marker Rab7. Equal amounts of WT, N127143Q, or Rem-T7 expression constructs were transfected into mouse mammary HC11 cells. Rem-CT was detected using a T7-specific antibody. Appropriate Alex Fluor 488 (green color) and Alex Fluor 594 (red color) secondary antibodies were used for T7 and Rab7, respectively. DAPI (blue) stained the nucleus. Circles mark the colocalization of Rem-CT with Rab7. (B) Quantitation of Rem-CT colocalization with Rab7. Dots represent the colocalization coefficients of individual cells. The middle line in each group represents the median value of the group. The medians were 0.04 for both the wild-type and glycosylation mutants. The highest and lowest lines represent the interquartile range. The numbers of cells analyzed were 35 for the wild-type Rem-CT, and 30 for the N127143Q mutant. Statistical analysis was performed using nonparametric Mann-Whitney tests.

### Rem-CT levels are rescued after BFA treatment in the presence of DN p97/VCP

The Arf guanine nucleotide exchange factors (GEFs) release GDP from Arf proteins, leading to their activation (85-88). Arf1 activation has multiple effects on the cellular vesicular network, including formation of COPI vesicles that mediate retrograde trafficking from the Golgi to the ER, changes in the activity of lipid-modifying enzymes, and altered composition of the Golgi, such as loss of golgin160 (89). BFA treatment of mammalian cells promotes an inactive GEF-Arf1-GDP complex (90-92). Therefore, we tested whether an HA-tagged DN Arf1 (T31N) has the same effect as BFA on Rem-CT levels. We co-transfected expression vectors for the DN Arf1 and either wild-type or N127143Q mutant Rem-T7 into 293T cells. Cell lysates subjected to Western blotting with HA-specific antibody confirmed the expression of the DN Arf1 (Fig. 7A). Similar to BFA treatment, we observed reduced levels of wild-type Rem-CT and no effect on the levels of the Rem-CT glycosylation mutant (Fig. 7A, compare lanes 1 and 2 versus lanes 3 and 4). Equal loading of cellular extracts was verified by Western blotting with GAPDH-specific antibodies. Together with previous data, these results suggest that reduction of Rem-CT levels by BFA is mediated through Arf1.

**FIG 7.**
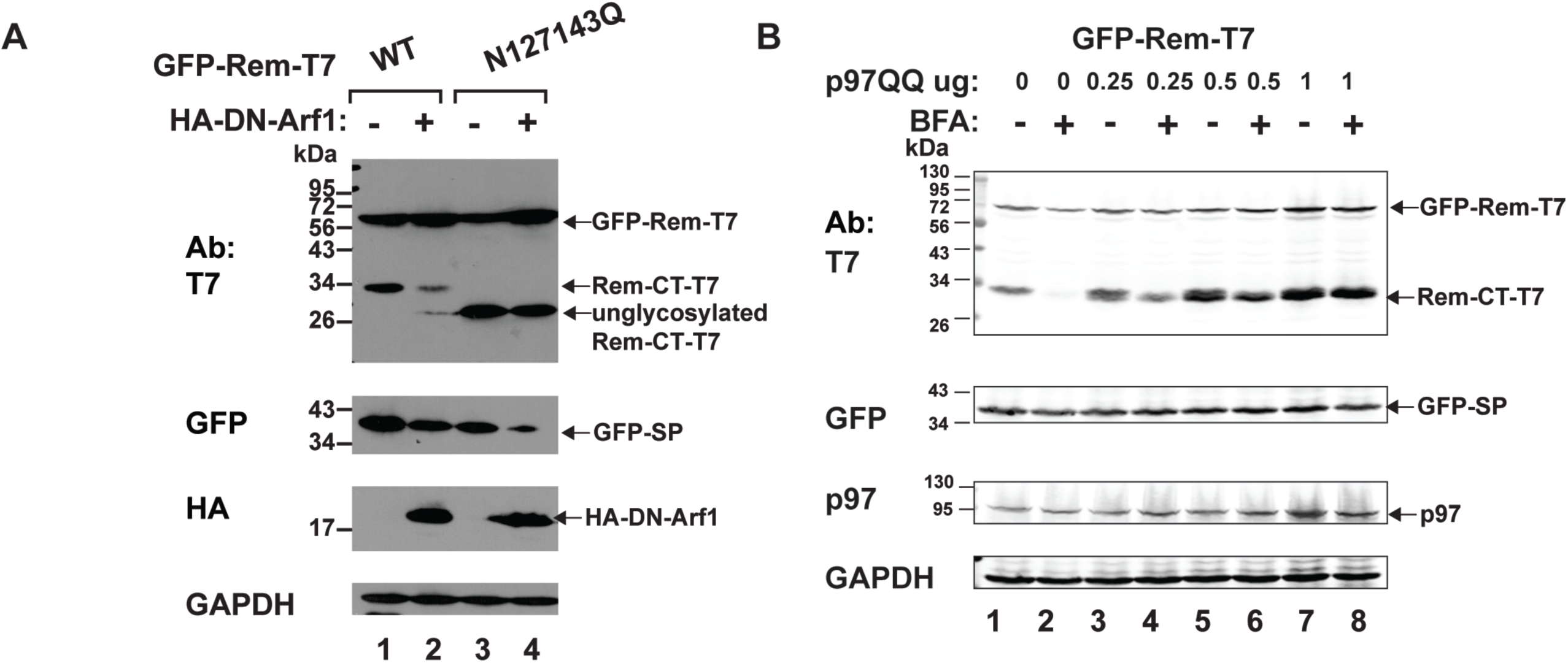
Dominant-negative p97/VCP rescues Rem-CT levels in the presence of BFA. (A) Expression of DN Arf1 mimics the reduced Rem-CT levels observed after BFA treatment. Cells (293T) were co-transfected with equal amounts of expression vectors for wild-type GFP-Rem-T7 (lanes 1 and 2) or the N127143Q glycosylation mutant (lanes 3 and 4) and DN Arf1. Cells were lysed and subjected to Western blotting with T7, GFP, HA, or GAPDH-specific antibodies. Arrows indicate the positions of tagged Rem precursor, glycosylated Rem-CT, or unglycosylated Rem-CT. (B) A DN form of p97 rescues Rem-CT levels in the presence of BFA. Increasing amounts of an expression plasmid for DN p97 (p97QQ) was transfected together with a constant amount of the GFP-Rem-T7 expression construct. Cells were treated with 3 μg/ml of BFA as indicated. Cells were lysed and subjected to Western blotting with T7, GFP, p97, or GAPDH-specific antibodies. LI-COR imaging was performed after incubation with appropriate secondary antibodies.

Our previous experiments indicated that Rem stability is regulated by the AAA ATPase p97/VCP (7, 8). In addition, SP extraction from the ER membrane for nuclear function is dependent on p97. Further, p97 has a role in regulation of vesicle fusion during reformation of the ER and Golgi after mitosis (93-95) as well as receptor-mediated endocytosis (96-99). Therefore, we determined whether p97 also affected the response of Rem-CT to BFA. Cells were co-transfected with a GFP-Rem-T7-expression vector and increasing amounts of a plasmid expressing the DN p97 (p97QQ) (100) in the presence or absence of BFA. Lysates were subjected to Western blotting with GAPDH-specific antibodies to confirm equal loading of extracts (Fig. 7B). Use of p97-specific antibodies revealed increased levels of the ATPase after transfection with the DN p97 expression plasmid. Examination of SP levels revealed that DN p97 did not affect Rem cleavage, consistent with our previously reported results (8). As expected, incubation of Western blots with T7-specific antibodies confirmed reduced levels of Rem-CT in the presence of BFA. However, expression of increasing amounts of the DN p97 completely reversed the BFA effect (Fig. 7B, compare lanes 2 and 8). These results suggest that DN p97 prevents Rem-CT trafficking to a cellular compartment promoted by treatment with BFA or expression of DN Arf1.

## DISCUSSION

We previously have shown that Rem is a truncated version of the MMTV Env protein with unusual trafficking and functional properties (2, 4, 7-10). Rem is a precursor protein that is directed for translation to the ER membrane where signal peptidase generates SP and the glycosylated Rem C-terminal product (Rem-CT) (Fig. 1A) (7, 9, 10). SP is then retrotranslocated from the ER membrane to the cytosol using the p97 ATPase (8, 9). Nuclear import of SP allows binding to the cis-acting Rem-responsive element (RmRE) found in all known MMTV mRNAs to facilitate nuclear export and post-export functions (2-4, 8, 9). Our recent studies show that loss of Rem C-terminal sequences antagonize the mutagenic effects of Apobec cytidine deaminases, particularly by proteasomal degradation of activation-induced cytidine deaminase (AID) during MMTV replication in lymphocytes (10). Since Apobec proteins are believed to function in both the cytoplasm and nucleus (101), determination of Rem-CT cellular localization was key to understanding its role in Apobec antagonism. In this paper, we showed that despite lacking a C-terminal KDEL/KKXX sequence (33, 35) and transmembrane domains, the steady state of Rem-CT remains in the ER. Furthermore, our data demonstrated that Rem-CT, like SP, has an unusual trafficking scheme that allows a small portion of the protein to localize to early and late endosomes. Trafficking of Rem-CT to early endosomes, but not late endosomes, appears to be prevented by lack of glycosylation, but does not involve passage through Golgi compartments that allow addition of complex glycans (59, 60) and requires p97/VCP. To our knowledge, no other viral or host proteins have been described with such a trafficking pathway.

A primary indicator of this unique pathway was our observation that BFA, a fungal metabolite that disrupts intracellular membrane trafficking between the ER and Golgi (49-54), reduced Rem-CT levels (Figs. 1B, 2, 4A, 4B, 7, S1, S2, and S4). We identified several factors that affected the sensitivity of Rem-CT to BFA. First, elimination of glycosylation sites either through substitution or deletion mutation prevented reduction of Rem-CT levels in the presence of BFA (Figs. 2, 4B, and S1). Our results indicated that glycosylated Rem-CT contains only simple glycans as determined by sensitivity to Endo H cleavage (Fig. 1D). Thus, Rem-CT does not traffic through the medial Golgi where modification to complex glycans occurs (59, 60). Second, co-expression of Rem-CT and a DN form of Arf1 mimics the effects of BFA (Fig. 7A). Arf1 participates in multiple parts of the secretory pathway, and the GBF and BIG subfamilies of Arf GEFs are affected in the presence of BFA by stabilizing the inactive GEF-ARF1-GDP complex (89). Moreover, Arf1 contributes to Golgi disassembly, particularly during M phase of the cell cycle (93, 95). Third, we cannot rescue Rem-CT in the presence of BFA using known inhibitors of degradation pathways, such as the proteasome or lysosome (Fig. 2), and Rem-CT is not detectably secreted in the presence or absence of the drug (Figs. 1B and S2). In addition, inhibitors of caspases, serine proteases, cathepsins, calpains, and metalloproteases fail to rescue Rem-CT from BFA treatment (data not shown). Fourth, Rem transcription and translation are not reduced by this fungal metabolite since the Rem precursor and SP expression are unaffected (Figs. 1B, 2, 4A, 4B, 7B, S1, and S4). Fifth, a DN form of p97/VCP prevents reduction of Rem-CT levels in the presence of BFA (Fig. 7B). Since p97 is involved in multiple cellular processes that involve assembly and disassembly of large protein complexes as well as membrane or vesicle fusions, we propose that Rem-CT has a function that requires trafficking outside the ER.

To our knowledge, Rem-CT is the only reported viral protein that displays this type of BFA sensitivity. Previous data suggested that two other cellular proteins, caspase 3 (CASP3) and Golgi-localized endoplasmic reticulum α-1, 2-mannosidase (ER ManI) have lower levels in the presence of BFA (102, 103). However, caspase 3 levels were rescued by a proteasome inhibitor, and this effect was observed only in one ovarian cell line W1PR (102). This situation differs from Rem-CT since the proteasome inhibitor MG132 did not restore Rem-CT levels in the presence of BFA (Fig. 2A). Moreover, the BFA sensitivity of Rem-CT appears to involve a conserved pathway since we observed this effect in human embryonic kidney as well as mouse mammary epithelial cells (Figs. 1B, 2, and S1). ER-ManI in the medial-Golgi binds the gamma subunit of COP-I (104) to mediate retrieval of ERAD substrates from the Golgi to the ER (103). Nevertheless, this study did not report a mechanism for reduced ER-ManI levels in the presence of BFA.

Confocal microscopy and immunostaining as well as cell fractionation were used to determine the cellular location of Rem-CT (Figs. 1C, 3, 5, and 6). We observed that the majority of wild-type Rem-CT was in the membrane fraction, and both the wild-type protein and the N127143 glycosylation mutant colocalized with the ER marker (Fig. 3A-C). However, a small portion of wild-type Rem-CT traffics to the early and late endosomes (Figs. 5 and 6). Both the wild-type protein and glycosylation mutant trafficked to the late endosomal compartment as determined by colocalization with the Rab7 marker (Fig. 6). In contrast, N127143Q showed significantly less colocalization with the early endosomal marker (Rab5) compared to wild-type Rem-CT (Fig. 5). The ER and endosomes have numerous contact sites for cholesterol transfer, endosome positioning, trans-Golgi to endosome protein transport, as well as receptor dephosphorylation and endosome fission (105-112). It is possible that trafficking through the endosomal system allows Rem-CT to counteract cellular proteins that function in host antiviral defenses. For example, tetherin/BST2, localizes to the cell surface and endosomes through the classical ER to Golgi transport system to inhibit viral release. The HIV-1-encoded Vpu protein reverses this inhibition by redirecting BST-2 from the cell surface to late endosomes (12, 13, 15, 18, 19, 22-25). Moreover, MHC-I, which is exported through the classical ER-Golgi pathway to the cell surface, is sequestered by HIV-1-encoded Nef within early endosomes. Redirection of MHC-I by Nef into late endosomes allows HIV-1 to evade immune responses (113).

We also attempted to map sequences that were required for the unique Rem-CT trafficking scheme. Rem-CT does not have a transmembrane domain or a canonical KDEL/KKXX at its C terminus, but the lack of a retrograde signal for the ER may be explained by its failure to proceed through the canonical secretory pathway. We performed deletion analysis and motif searches (Fig. 4 and S4), but no specific sequence emerged that provided a retrograde trafficking signal. It is possible that the retrograde signal is redundant, thus preventing detection. Alternatively, recent evidence from plant cell experiments indicates that BFA relocalizes a number of Golgi enzymes to the ER. Specific enzymes typical of the cis-Golgi traffic, using a COPII-independent mechanism, enter a compartment known as the Golgi entry core compartment (GECCO) near ER exit sites (ERES) in the presence of BFA (104). GECCO may be similar to the ERGIC, which serves as a sorting station for proteins at ERES (114-117). We propose that Rem-CT leaves the ERES and traffics directly to late endosomes or indirectly to the ERGIC prior to endosomal localization (Fig. 8). Rem-CT sorting to early endosomes and the ERGIC is dependent on glycosylation and, perhaps, p97/VCP instead of COPII. BFA treatment, which collapses the Golgi cisternae into the ERGIC, may lead to Rem-CT proteolysis. This model leaves many unanswered questions, but suggests that uncleaved Rem, rather than Rem-CT, may be the antagonist of Apobec family enzymes (10).

**FIG 8.**
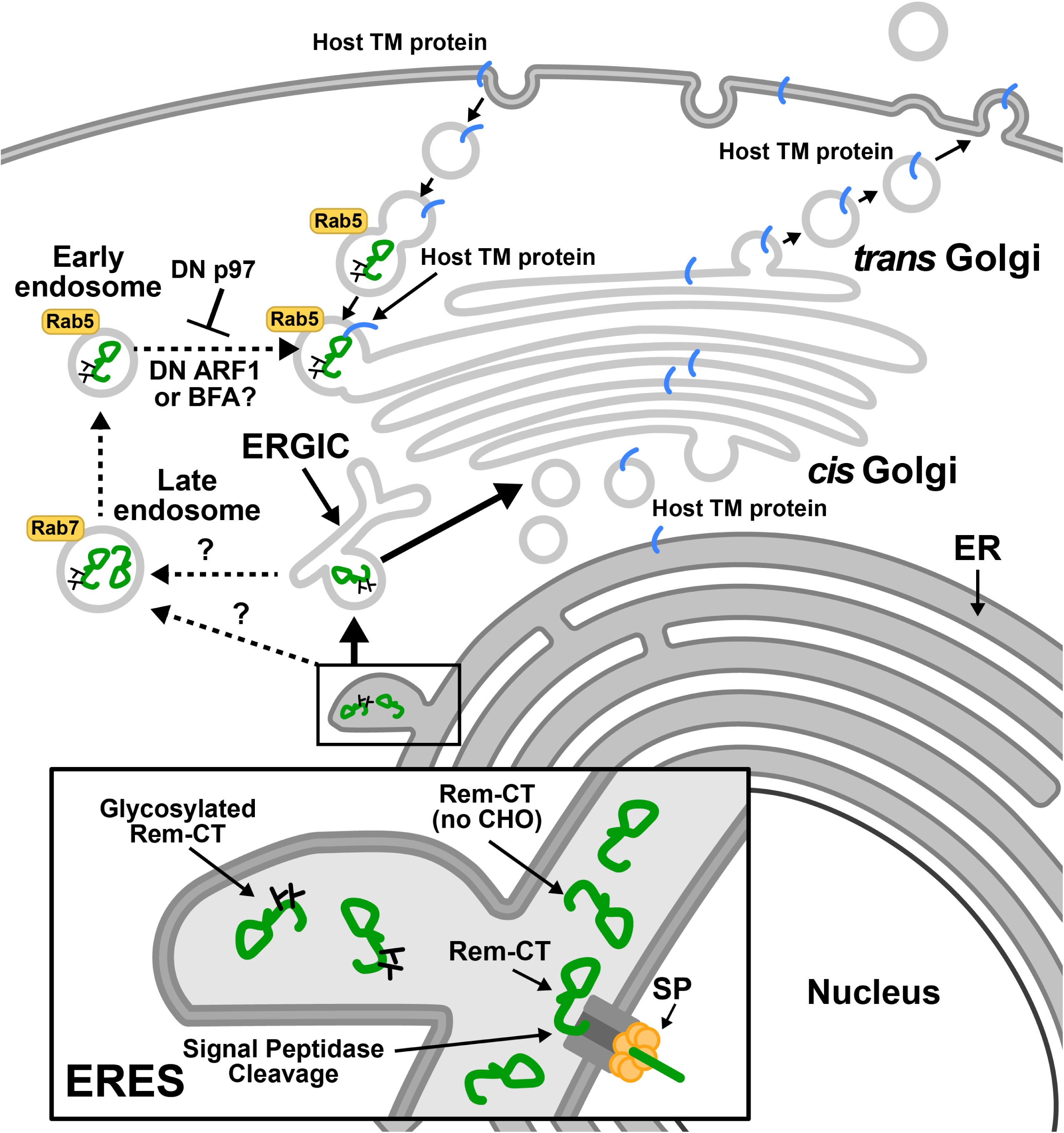
Model for Rem-CT trafficking. Rem is directed for translation to the ER membrane and cleaved by signal peptidase into an N-terminal SP and Rem-CT. SP is retrotranslocated into the cytosol by the p97 ATPase and traffics to the nucleus to perform a Rev-like function. The Rem-CT cleavage product is glycosylated with simple glycans and maintains high steady state levels in the ER lumen. Glycosylation of Rem-CT at one of two potential sites is required for trafficking out of the ER. A portion of both glycosylated and unglycosylated Rem-CT exits the ER, presumably at ERES, to the late endosomes. Only glycosylated Rem-CT is detectable in early endosomes. BFA treatment and expression of DN Arf1 lead to reduction of Rem-CT levels, whereas DN p97 ATPase restores those levels, perhaps by blocking vesicle formation. Rem (no CHO) = Rem lacking glycosylation (carbohydrate).

In summary, our study identifies the first viral protein that uses this unique BFA-sensitive pathway. The benefit of this trafficking for viruses and host cell proteins requires additional experiments. Nevertheless, the study of viruses continue to elucidate novel cell processes that may allow development of new antiviral and cancer therapies.

## MATERIALS AND METHODS

### Cell culture and transfection

HEK293 or 293T cells were cultured in Dulbecco’s modified Eagle’s medium (Sigma-Aldrich) supplemented with 10% fetal calf serum, gentamicin sulfate (50 μg/ml), penicillin (100 units/ml), and streptomycin (50 μg/ml). HC11 cells were maintained in RPMI media (Sigma-Aldrich) supplemented with 10% fetal calf serum, gentamicin sulfate (50 μg/ml), insulin (0.5 μg/ml), and EGF (1 μg/ml). Cells were routinely tested for contamination with bacteria using a PCR-based assay. Transfections of HEK293 or 293T cells were performed using polyethylenimine (PEI) (Polysciences) transfection. Briefly, 1 x 10^6^ cells were seeded into each well of a 6-well plate on the day prior to transfection. DNA (3 μg) was mixed with 200 μl of DMEM and 9 μl of PEI (1 μg/μl) and incubated at room temperature for 20 min before adding to pre-seeded cells. At 7 h post-transfection, cells were washed with phosphate-buffered saline (PBS) once and replaced with fresh media. Transfections of HC11 cells were performed using Lipofectamine 3000 (Thermo Fisher Scientific) using protocols from the manufacturer. Cells were harvested 48 h after transfection. For some experiments, Brefeldin A (Cell Signaling Technology) or MG132 (Boston Biochem) or Bafilomycin A1 (Sigma-Aldrich) were added at the indicated concentrations for 6 h before cells were harvested. Endoglycosidase H (Endo H, NEB P0702S) (500 U) was added to 20 μg of cell extracts for 1 h at 37°C prior to analysis by Western blotting.

### Plasmid constructions and sources

GFP-Rem-T7 and Rem-T7 plasmids were constructed based on previously described N-terminally GFP-tagged (GFP-Rem) or Rem (3) with the addition of six glycine linkers and the T7 tag sequence (MASMTGGQQMG) at the C-terminus as indicated. CMV-SEAP was a gift from Alan Cochrane (Addgene plasmid #24595; http://n2t.net/addgene:24595; RRID:Addgene_24595). The CMV-SEAP-T7 construct was made based on plasmid CMV-SEAP with the addition of six glycine linkers and the T7 tag at the C-terminus. Linkers and the T7 tag were added using PCR and the CloneAmp HiFi PCR Premix (TakaRa). Mutants (N127143Q and motif substitutions) were obtained by mutagenesis PCR using CloneAmp HiFi PCR Premix (TakaRa). The GFP-tagged N127143Q mutant was constructed by substitution of asparagines for glutamines at amino acids 127 and 143 by site-directed mutagenesis (Stratagene). The GFP-Rem-T7 partial deletion mutants (GFP-RemΔ103-155-T7, GFP-RemΔ155-198-T7, GFP-RemΔ198-247-T7, and GFP-RemΔ247-30-T7) were constructed similarly from GFP-Rem-T7 by respectively deleting the designated 50 amino acid segments from the Rem C-terminus. These mutations were prepared by inserting *SalI* sites into the *rem* gene by site-directed mutagenesis at the designated codon sites followed by digestion with *SalI* and religation of the vector. Forward and reverse primers were used for PCR in separate tubes with the following parameters: 98°C for 1 min, 5 cycles at 98°C for 10 s, 55°C for 30s, and 72°C for 3.5 min (or based on the final product length, 0.5 min per kb) with a final extension at 72°C for 30 min. Subsequently, the forward and reverse primer PCRs were mixed and used for additional cycles: 98°C for 1 min, followed by 20 cycles at 98°C for 10 s, 55°C for 30s, and 72°C for 3.5 min (or based on the final product length, 0.5 min per kb) with final extension at 72°C for 30 min. All constructs were sequenced to confirm the mutations, and restriction digests were used to confirm the plasmid integrity.

The ER-mCherry expression plasmid was a gift from Dr. N. Hosokawa (Kyoto University, Kyoto, Japan) (76). The HA Arf1 DN (T31N) plasmid was a gift from Thomas Roberts (Addgene plasmid # 10833; http://n2t.net/addgene:10833; RRID: Addgene_10833) (118). The CFP-VSV-G construct was a gift from Michael Davidson (Addgene plasmid # 55397; http://n2t.net/addgene:55397; RRID: Addgene_55397).

### Luciferase reporter assays

The SP-responsive reporter assays have been described previously (7-9) using the pHM*Rluc* Rem-responsive construct (4). The pHM*Rluc* plasmid contains the 3’ end of the C3H-MMTV genome missing the MMTV SP coding sequence. Insertion of the *Renilla* luciferase reporter gene between splice donor and acceptor sites within the MMTV envelope gene avoided interruption of the Rem-responsive element (4). Briefly, transfections were performed in triplicate by co-transfection of the pHM*Rluc* Rem-responsive construct and a Rem-non-responsive firefly luciferase reporter construct (pGL3-Control; Promega), in the presence and absence of a Rem-expression plasmid. After 48 h, transfected cells were lysed in buffers provided with the Dual Luciferase assay kit (Promega). Protein concentrations were determined (Bio-Rad Protein Assay System), and lysates (∼40 μg) were used to detect firefly and *Renilla* luciferase activity. Rem-responsive *Renilla* luciferase data were then normalized to Rem-non-responsive firefly luciferase levels to control for differences in transfection efficiency. Values are reported for 100 μg of protein lysate. The means of normalized *Renilla* luciferase activity for triplicate transfections with standard deviations are shown.

### Western blotting

Western blots were performed as previously described (8). Antibodies were obtained from sources as described in parentheses: monoclonal T7 (Cell Signaling Technology), GFP (Clontech), GAPDH (Cell Signaling Technology), p97/VCP (Invitrogen), and LC-3B (Cell Signaling Technology). Horseradish peroxidase-conjugated secondary antibodies were obtained from Cell Signaling Technology. Fluorescently labeled IRDye secondary antibodies were purchased from LI-COR Biosciences.

### Cell supernatant concentration

After cells were treated with BFA, the medium was collected and subjected to centrifugation to remove cell debris. Supernatants were filtered through a 0.45 micron filter. Concentration of supernatants was performed based on the manufacturer’s instructions. Briefly, Amicon® Ultra-15 Centrifugal Filter Units were rinsed with Milli-Q water before use, and then the filtered supernatants were added into each unit. Filter units were used for centrifugation at 4,000 Xg for 40 min, and the concentrated solute was recovered from the unit for Western blotting analysis.

### Exosome isolation

After cells were treated with BFA, the medium was collected and subjected to centrifugation to remove cell debris. Supernatants were filtered through a 0.45 micron filter. The miRCURY Exosome kits were purchased from Qiagen (Cat #76743), and exosome isolation was performed according to the manufacturer’s instructions. Briefly, 0.4 volume of Precipitation Buffer B was added for every 1 volume of the sample and mixed thoroughly. The precipitation was performed overnight at 4°C, followed by centrifugation and supernatant removal. Resuspension buffer was used for pellets, which were processed for Western blotting.

### Cellular fractionation

Transfected cells were harvested 48 hr post transfection. Cells were washed with PBS twice to remove the residual media, and then permeabilized for 5 min on ice with 0.02% (w/v) digitonin using a fractionation buffer containing Roche protease inhibitor cocktail. The cytosolic fraction was isolated from the cells by centrifugation at 13,000 Xg for 5 min at 4°C. The remaining pellet was incubated in 1% Triton X-100 for 5 min on ice, followed by a second centrifugation at 13,000Xg for 5 min at 4°C to separate the membrane and nuclear fractions.

### Immunofluorescence and image analysis

On the day of transfection, coverslips were washed with ethanol, placed into 6-well plates, and dried under UV light in a laminar flow hood. After 24 h, coverslips were washed with PBS once, and ∼0.6 × 10^6^ cells were trypsinized and plated in each well. Transfected cells were incubated for 48 h and fixed with 4% paraformaldehyde in PBS. Cells were permeabilized with 0.1% Triton-X-100 and then washed three times with PBS containing 0.1% Tween-20 (PBST) for 5 min each. Cells were then blocked with blocking buffer (2% FBS, 0.1% Tween-20, 150 mM NaCl, 10 mM Tris-HCl, pH 7.4) for 1 h at room temperature and then incubated with primary antibodies overnight at 4°C. After three PBST washes following the primary antibody incubation, Alexa Fluor secondary antibodies (Invitrogen, Thermo Fisher Scientific) were added for 2 h at room temperature. Samples were then washed thrice with PBST and stained with DAPI (150 nM) for 10 min at room temperature. After three PBST washes, coverslips were mounted on slides with VECTASHIELD Antifade Mounting Media (Vector Laboratories) and sealed for fluorescence imaging. Monoclonal Rab5- and Rab7-specific antibodies were purchased from Cell Signaling Technology. When two different primary antibodies were used, the first primary antibody was incubated overnight before samples were washed with PBST three times. The second primary antibody then was incubated with the sample overnight. During secondary antibody incubation, Alexa Fluor secondary antibodies with different spectral characteristics from different animal sources were used to ensure recognition of only one primary antibody.

The Zeiss LSM 710 Confocal and Elyra S.1 Structured Illumination Super Resolution Microscope was used to capture Z-stack sample images. All slices in a Z stack across samples were kept at 0.33 μm per slice interval. Lasers (405nm, 488nm and 561nm) were used with the same power and gain across samples to obtain images. Quantitation and image processing were performed using Zeiss software ZEN, Graphpad (Prism) and ImageJ. Briefly, the Mander’s Colocalization Coefficient (MCC) (119) was used to quantify the fraction of Rem-CT colocalized with ER-mCherry or other markers (Rab5/Rab7) using pixels from each cell in single slices and ZEN software. Mann-Whitney’s t tests were used to compare different samples in Graphpad. Images were processed in ImageJ.

### Reproducibility and statistical analysis

All experiments were performed two to five times with similar results. Individual statistical tests are indicated in each section or figure. A p-value of <0.05 was considered to be significant.

## ACKNOWLEGMENTS

We thank members of the Dudley laboratory for helpful comments on experimental design and the manuscript. We also are grateful for advice from the Microscopy and Imaging Core facilities at The University of Texas at Austin. This work was supported by National Institutes of Health grant R01 AI131660.

---

**FIG S1** Reduced levels of Rem-CT after BFA treatment is prevented by lack of glycosylation in mouse mammary epithelial cells. HC11 cells were transfected with equal amounts of expression vectors for GFP-Rem-T7 (lanes 1 and 2) or dually tagged N127143Q (lanes 3 and 4) and treated with or without 3 μg/ml BFA. Cells were lysed and subjected to Western blotting with T7, GFP, or GAPDH-specific antibodies. LI-COR imaging was performed after incubation of Western blots with appropriate secondary antibodies. The asterisk indicates the unglycosylated Rem-CT protein produced by the mutant N127143Q (lanes 3 and 4).

**FIG S2**. Rem-CT is not detectable in exosomes after BFA treatment. Cells (293T) were transfected with expression plasmids for GFP-Rem-T7 and treated with or without 3 μg/ml BFA. Supernatants were filtered prior to isolation and concentration of exosomes as described in Materials and Methods. Cell lysates (lanes 1 and 2) and concentrated exosomes (lanes 3 and 4) were subjected to Western blotting using T7 or GAPDH-specific antibodies.

**FIG S3** Luciferase assays confirm SP function from Rem-T7 constructs. Cells (293T) were transfected with either 10 or 20 ng of Rem-T7 constructs together with luciferase reporter constructs as described in Materials and Methods. Luciferase activities normalized for transfection efficiency were reported as the means and standard deviations from triplicate transfections. The sample lacking a Rem expression construct (pcDNA3) was assigned a relative value of 1.

**FIG S4** Substitution mutants of putative trafficking motifs within Rem-CT fail to affect the response to BFA. (A) Potential cellular trafficking motifs were identified by the Eukaryotic Linear Motif (ELM) Prediction Tool within Rem-CT. The motifs are identified in separate colors as WELW (clathrin binding), YPHC (an SH3 interaction motif for cytoskeletal organization and membrane trafficking), and PPKY (a WW domain recognition motif for protein interactions). The introduced mutations are shown below the wild-type sequence. (B), (C) and (D) show the results of transfections of 293T cells with expression constructs for wild-type GFP-Rem-T7 or the same amounts of the glycosylation mutant N127143Q or motif mutants. Transfected cells were treated with or without 3 μg/ml BFA. Cells were lysed and subjected to Western blotting with T7, GFP or GAPDH-specific antibodies. The YPHC mutant produced no detectable GFP-tagged SP.

